# Physicochemical mechanotransduction alters nuclear shape and mechanics via heterochromatin formation

**DOI:** 10.1101/423442

**Authors:** Andrew D. Stephens, Patrick Z. Liu, Viswajit Kandula, Haimei Chen, Luay M. Almassalha, Vadim Backman, Thomas O’Halloran, Stephen A. Adam, Robert D. Goldman, Edward J. Banigan, John F. Marko

## Abstract

The nucleus houses, organizes, and protects chromatin to ensure genome integrity and proper gene expression, but how the nucleus adapts mechanically to changes in the extracellular environment is poorly understood. Recent studies have revealed that extracellular chemical or physical stresses induce chromatin compaction via mechanotransductive processes. We report that increased extracellular multivalent cations lead to increased heterochromatin levels through mechanosensitive ion channels. This increase in heterochromatin results in increased chromatin-based nuclear rigidity, which suppresses nuclear blebbing in cells with perturbed chromatin or lamins. Furthermore, transduction of elevated extracellular cations rescues nuclear morphology in model and patient cells of human diseases, including progeria and the breast cancer model cell line MDA-MB-231. We conclude that nuclear mechanics and morphology, including abnormal phenotypes found in human diseases, can be modulated by cell sensing of the extracellular environment and consequent changes to histone modification state and chromatin-based nuclear rigidity, without requiring direct mechanical perturbations to the cell interior.

## Introduction

The nucleus is the organelle within the cell that contains, organizes, and mechanically protects the genome. Nuclear mechanics dictates proper genome organization and deformations, both of which can alter gene expression (Cremer and Cremer, 2001; Tajik et al., 2016; Cho et al., 2017; Kirby and Lammerding, 2018). Consequently, abnormal nuclear morphology and mechanics occur across a spectrum of human diseases and conditions, including heart disease, muscular dystrophy, aging, and many cancers (Butin-Israeli et al., 2012). These conditions are linked to perturbations of the major mechanical components of the nucleus, chromatin and lamins, and they result in nuclear rupture and abnormal deformations termed “blebs” (Goldman et al., 2004; Vargas et al., 2012; Furusawa et al., 2015; Robijns et al., 2016; Stephens et al., 2018). Cellular and extracellular compressive forces, such as those due to actin and migration though tight spaces further exacerbate nuclear morphological instability (Khatau et al., 2009; Tamiello et al., 2013; Denais et al., 2016; Raab et al., 2016; Hatch and Hetzer, 2016; Tocco et al., 2017). Subsequently, abnormal nuclear shape and rupture lead to mislocalization of nuclear proteins, DNA damage, and altered transcription, all of which are thought to cause cellular dysfunction in human diseases (Shimi et al., 2008; Isermann and Lammerding, 2013; Pfeifer et al., 2018; Xia et al., 2018). It is unclear how the cell natively regulates nuclear mechanics and morphology to prevent these outcomes when subject to mechanical stresses.

Treatment of cells with a broad histone demethylase inhibitor to increase heterochromatin and nuclear rigidity has been shown to rescue nuclear shape in cases of abnormal morphology (Stephens et al., 2018). Alterations to chromatin compaction on both short and long time scales can be achieved through changes in extracellular media or forces applied to the cell, as shown in studies that varied osmolality (Albiez et al., 2006; Irianto et al., 2013), cell substrate stretching (Heo et al., 2015; Le et al., 2016; Heo et al., 2016; Gilbert et al., 2018 Preprint), substrate patterning (Versaevel et al., 2012; Jain et al., 2013), and cell migration (Gerlitz and Bustin, 2010; Jacobson et al., 2018 Preprint; Segal et al., 2018 Preprint). Direct loading by stretching the underlying cell substrate results in chromatin compaction that depends on the heterochromatin methyltransferase EZH2 (Heo et al., 2015) and/or increases in the facultative heterochromatin marker H3K27me^3^ (Le et al., 2016; Heo et al., 2016). This pathway is dependent on mechanosensitive ion channels in the plasma membrane, which respond to membrane tension and stretching (Kim et al., 2015; Heo et al., 2015), and it can be blocked by inhibitors GdCl_3_ or tarantula venom GsMTx4 (Suchyna et al., 2000; Ermakov et al., 2010). This heterochromatin-forming mechanotransduction pathway is a candidate for native regulation of nuclear mechanics and morphology.

Prior studies have focused on mechanically stressing the entire cell to increase heterochromatin but have not fully investigated the effects on physical properties of the cell nucleus. We sought to determine whether activation of mechanosensitive ion channels, without whole-cell stretching, could trigger increases in heterochromatin that could regulate nuclear morphology via increased rigidity. Multivalent cations in the extracellular environment interact with and distort negatively charged phospholipids, thus increasing membrane tension and bending (Gleisner et al., 2016; Ali Doosti et al., 2017). We observed that adding divalent cations (MgCl_2_ and CaCl_2_) or cationic polyamines (spermidine) to normal cell culture media increased heterochromatin levels via mechanosensitive ion channels, while internal cell ion contents remained unchanged. Remarkably, the cation treatments rescued nuclear shape in cells displaying aberrant morphology (such as nuclear blebs) that had been induced by direct perturbation of either chromatin or lamin B. These cation-induced increases in heterochromatin were dependent on histone methyltransferases, and they led to increases in nuclear rigidity. Furthermore, supplementing media with increased cations similarly increased heterochromatin and rescued nuclear morphology in models of the accelerated aging disease Hutchison-Gilford progeria syndrome (HGPS) and breast cancer, as well as in patient HGPS cells. Our results show that the cell can modulate nuclear mechanics and morphology in response to the extracellular environment through mechanosensitive ion channels that cue changes in histone modification state, driving heterochromatin formation and increased chromatin-based nuclear rigidity.

## Results and Discussion

### Increased extracellular divalent cations increase heterochromatin levels through mechanosensitive ion channels

Previous work has revealed that cell stretching activates mechanosensitive ion channels, leading to increased heterochromatin (Heo et al., 2015; Le et al., 2016; Heo et al., 2016). To activate mechanosensitive ion channels without physically stretching the cell, we increased extracellular divalent ion concentrations; this is known to increase membrane tension (Gleisner et al., 2016; Ali Doosti et al., 2017). Incubation of mouse embryonic fibroblast (MEF) cells in increased extracellular levels of divalent cations (7.5 or 17.5 mM MgCl_2_ or CaCl_2_) resulted in increased facultative (H3K27me^3^) and constitutive (H3K9me^2,3^) heterochromatin, as measured by immunofluorescence and Western blots (Figure 1A-E, 24-hour treatment). Treatment with mechanosensitive ion channel inhibitors blocked increases in heterochromatin (GsMTx4 and GdCl_3_, Figure 1, C and F), similar to its effect in cell stretching experiments (Heo et al., 2015; Heo et al., 2016). Addition of divalent cations to the medium did not alter cell viability, growth, nucleus size, or actin content (Supplemental Figure 1). Thus, mechanosensitive ion channels activated by extracellular divalent cations can induce higher levels of heterochromatin independently of cell stretching.

**Figure 1.**
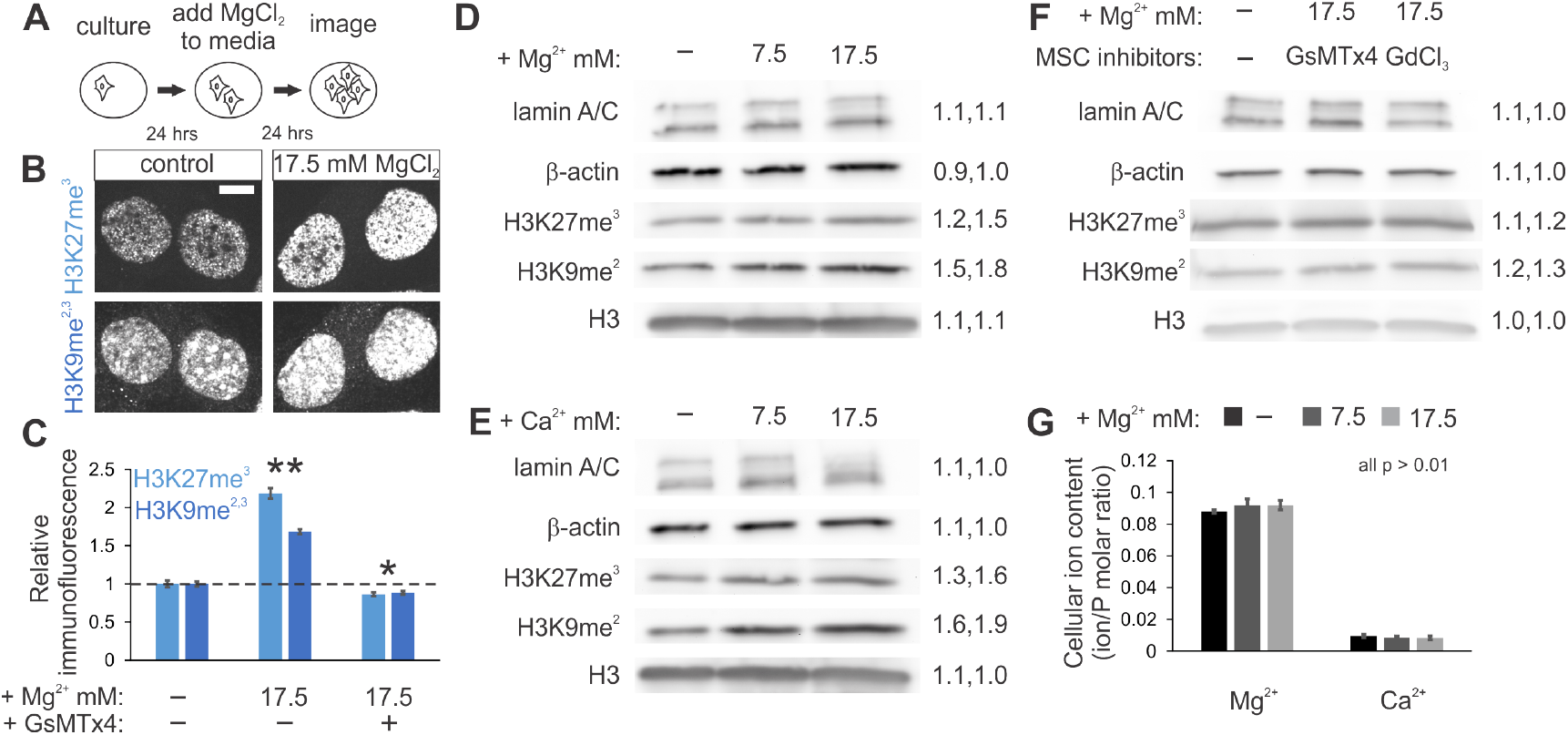
Increased extracellular divalent cations increase heterochromatin formation through mechanosensitive ion channels. (A) Schematic of MEF cell treatment for 24 hours, (B) representative images, and (C) average immunofluorescence signals for heterochromatin (H3K27me^3^ and H3K9me^2,3^) for cells incubated with or without additional MgCl_2_, with or without GsMTx4 (n = 54, 53, and 109, respectively). (D-F) Western blots (with quantifications, right) from cells in standard medium or supplemented with (D) MgCl_2_, (E) CaCl_2_, or (F) MgCl_2_ and mechanosensitive ion channel inhibitors (GsMTx4 and GdCl_3_) for 24 hours. (G) ICP-MS analysis of cellular Mg^2+^ and Ca^2+^ ion contents relative to phosphorous (P) content (n = 8, 8, and 5, respectively). Scale bar = 10 µm. Error bars represent standard error. Asterisks denote statistically significant differences from control (p < 0.01), where different numbers of asterisks are also significantly different.

To determine whether treatments that increased extracellular cations impacted intracellular levels of those cations, we performed inductively coupled plasma mass spectrometry analysis (ICP-MS), which quantifies levels of intracellular ions. ICP-MS analysis measured no change in intracellular Mg^2+^ and Ca^2+^ relative to phosphorous (Figure 1G) in 24-hour MgCl_2_ extracellular-cation-treated cells relative to untreated cells. Therefore, the extracellular-cation-triggered heterochromatin increase is not due to an increase in intracellular divalent cations, and for instance, consequent physicochemical condensation of chromatin via divalent cations (*e.g.*, (Poirier et al., 2002; Pajerowski et al., 2007; Stephens et al., 2017a; Shimamoto et al., 2017)). Instead, our data suggest that extracellular divalent cations increase membrane tension (Gleisner et al., 2016; Ali Doosti et al., 2017), which activates tension-sensitive ion channels. This subsequently leads to increased heterochromatin formation, similar to experiments in which direct cell substrate stretching was observed to elevate heterochromatin levels (Le et al., 2016; Heo et al., 2016).

### Extracellular multivalent cation transduction suppresses nuclear blebbing

To determine the impact of mechanotransduction-based heterochromatin formation on nuclear morphology, we investigated the effects of added extracellular MgCl_2_ on cells with disrupted nuclear morphology. We induced abnormal nuclear ruptures and “bleb” deformations by treatment with histone deacetylase inhibitor valproic acid (VPA), which increased decompacted euchromatin (Figure 2A; (Stephens et al., 2018)). Previously, we showed that nuclear morphology could be rescued by histone demethylase inhibition-driven increases in heterochromatin (Stephens et al., 2018). Here, we instead increased levels of extracellular MgCl_2_ in VPA-treated cells. This resulted in an increase in heterochromatin, as measured by immunofluorescence and Western blots, and a corresponding decrease in nuclear blebbing (Figure 2B-E; Supplemental Figure 2A, CaCl_2_ and 2B, HT1080 MgCl_2_). VPA-treated cells displayed nuclear blebs in 17% of cells, while adding 2.5 mM extracellular MgCl_2_ decreased nuclear blebbing to ∼12%, 7.5 mM to ∼10%, and 17.5 mM to ∼6.5%, showing a trend toward the 3-4% blebbing observed in control cells (Figure 2E). Time-course analysis revealed that VPA-based nuclear blebbing is rescued coincident with an increase in heterochromatin after 8 hours of incubation in increased extracellular MgCl_2_, but not after 2 hours, while euchromatin remains constant (Supplemental Figure 2A-D). This effect is dependent on the mechanotransduction pathway described above, as inhibition of mechanosensitive ion channels via GsMTx4 or GdCl_3_ inhibited the ability of increased extracellular divalent cations to suppress chromatin-decompaction-induced nuclear blebs (Figure 2G). Calcium secondary messengers calcineurin and calmodulin were also essential to extracellular cation-based bleb suppression while major signaling kinases (Ca^2+^/calmodulin-dependent kinase II and myosin light chain kinase) and global transcription were dispensable (Supplemental Figure 2H). Thus, cellular mechanotransduction of extracellular cues by mechanosensitive ion channels can rescue nuclear morphology coincident with increasing heterochromatin formation.

**Figure 2.**
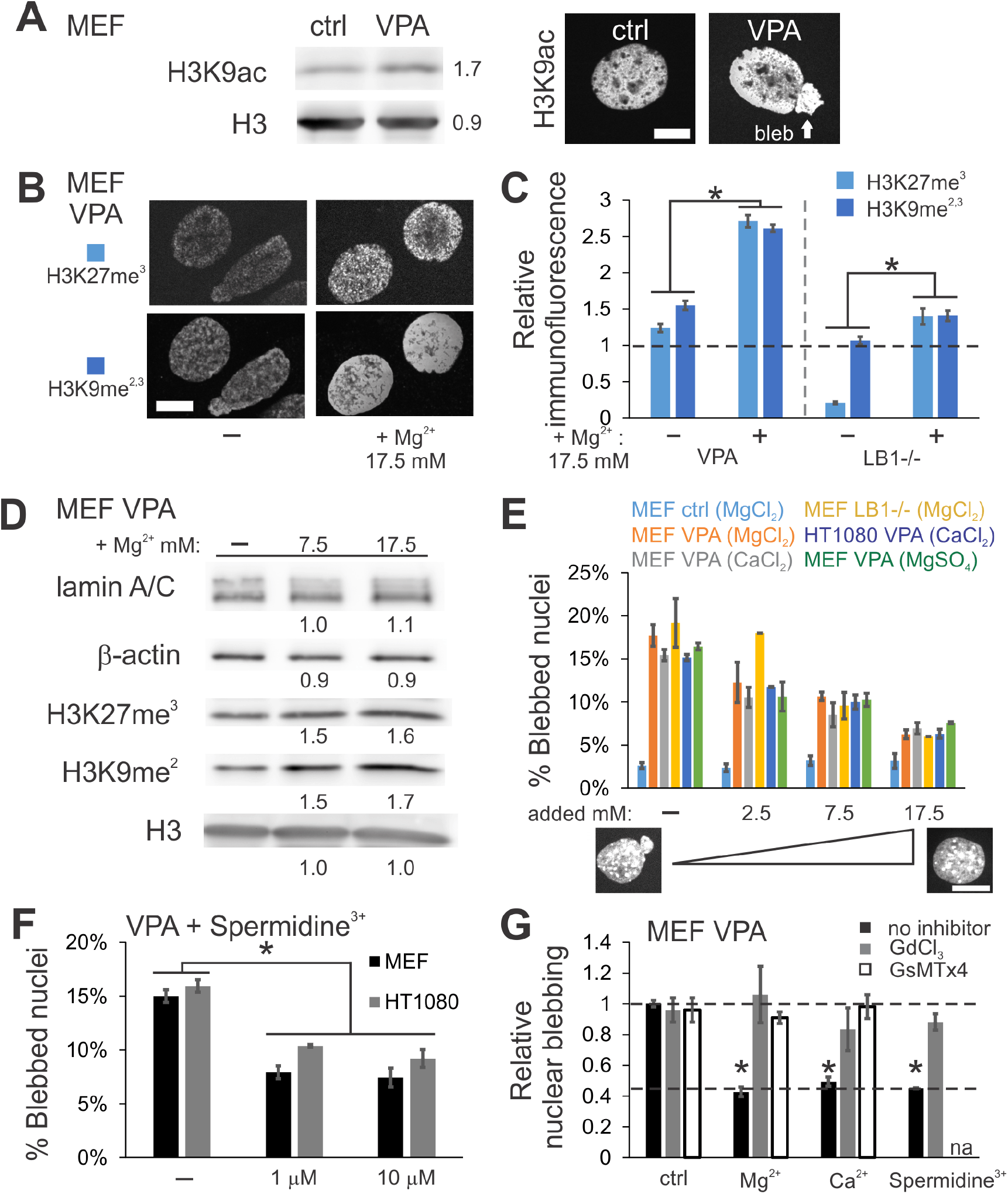
Increased extracellular divalent cations or cationic polyamines suppress nuclear blebbing. (A) Western blots and immunofluorescence images showing that valproic acid (VPA) increases euchromatin (H3K9ac) and nuclear blebbing relative to control (ctrl). (B) Representative images and (C) average heterochromatin (H3K27me^3^ and H3K9me^2,3^) immunofluorescence signals in VPA or lamin B null (LB1-/-) cells in media with normal and additional MgCl_2_ (n = 20 – 53, wild type scaled to 1). (D) Western blot from VPA-treated MEF cells in standard or MgCl_2_ supplemented medium for 24 hours. (E) Graph of percentage of nuclei displaying a nuclear bleb upon treatment with additional extracellular divalent cations or (F) spermidine^3+^ in the medium in both MEF and HT1080 cells (for IF see Supplemental Figure 2, E and F). (G) Graph of relative nuclear blebbing for VPA-treated cells with increased extracellular divalent cations and mechanosensitive ion channel inhibitors GsMTx4 and GdCl_3_ (VPA without inhibitor (ctrl) scaled to 1). Nuclear blebbing experiments, n = 3-5 experiments of 90-200 cells each. Scale bar = 10 µm. Error bars represent standard error. Asterisks denote statistically significant differences (p or χ^2^ < 0.05).

This native mechanotransduction response is general and physiologically relevant. An alternative cell model of nuclear rupture and blebbing is the lamin B1 null (LB1-/-) cell line (Vargas et al., 2012), which also has decreased heterochromatin. Similar to VPA-treated cells, LB1-/-cells exhibited increases in heterochromatin and decreased nuclear blebbing in the presence of additional extracellular divalent cations (Figure 2, C and E). Suppression of nuclear blebbing in perturbed cells was similar when using a different divalent cation (CaCl_2_), counterion (MgSO_4_), or cell type (HT1080) (Figure 2E). VPA-treated cells exposed to the physiologically relevant cationic polyamine spermidine^3+^ at physiological concentrations (µM) revealed a similar decrease in nuclear blebbing. Again, this effect was dependent on mechanosensitive ion channels and coincided with increased heterochromatin (Figure 2F-G, Supplemental Figure 2F). The low concentration used rules out an osmotic origin for the observed effects. This finding reveals that mechanotransduction-based heterochromatin formation and nuclear morphology rescue is general and can be activated by physiologically relevant molecules. Thus, the extracellular environment sensed by mechanosensitive channels in the cell membrane can influence nuclear morphology by increasing heterochromatin over hours, which in turn suppresses the biophysical effects of direct chromatin or lamin perturbations that cause abnormal morphology.

These results are significant because the ability of the nucleus to maintain its shape directly impacts genome stability and transcription. Nuclear blebbing coincides with nuclear rupture (Vargas et al., 2012; Tamiello et al., 2013; Robijns et al., 2016; Hatch and Hetzer, 2016; Stephens et al., 2018), which has been tied to DNA damage (Denais et al., 2016; Raab et al., 2016; Pfeifer et al., 2018; Xia et al., 2018). Furthermore, nuclear blebs can disrupt proper transcription, as chromatin within blebs exhibit decreased transcription (Shimi et al., 2008; Helfand et al., 2012; Bercht Pfleghaar et al., 2015). Therefore, activation of mechanosensitive ion channels serves as a native and tunable mechanism to mechanically regulate nuclear stability and function via heterochromatin formation.

### Transduction of extracellular cues to rescue nuclear morphology is dependent upon methyltransferases to increase heterochromatin levels

To determine whether rescue of nuclear morphology by extracellular divalent cations was dependent on the corresponding increases in heterochromatin, we co-treated cells with additional MgCl_2_ and the broad histone methyltransferase inhibitor DZNep (Miranda et al., 2009). DZNep-treated cells have decreased heterochromatin and display increased levels of nuclear blebbing similar to VPA (Figure 3A; (Stephens et al., 2018)). In stark contrast to control and VPA-treated cells, cells co-treated with DZNep and additional extracellular cations displayed no increase in heterochromatin markers, as measured by Western blots, and nuclear blebbing was not suppressed (Figure 3B-D). This is a key result because both histone deacetylase and methyltransferase inhibitors result in net chromatin decompaction, weakened chromatin-based nuclear rigidity, and increased nuclear blebbing. However, only the histone methyltransferase inhibitor prevents transduction of these extracellular environmental cues into nuclear morphological rescue, indicating that heterochromatin formation is essential to the process.

**Figure 3.**
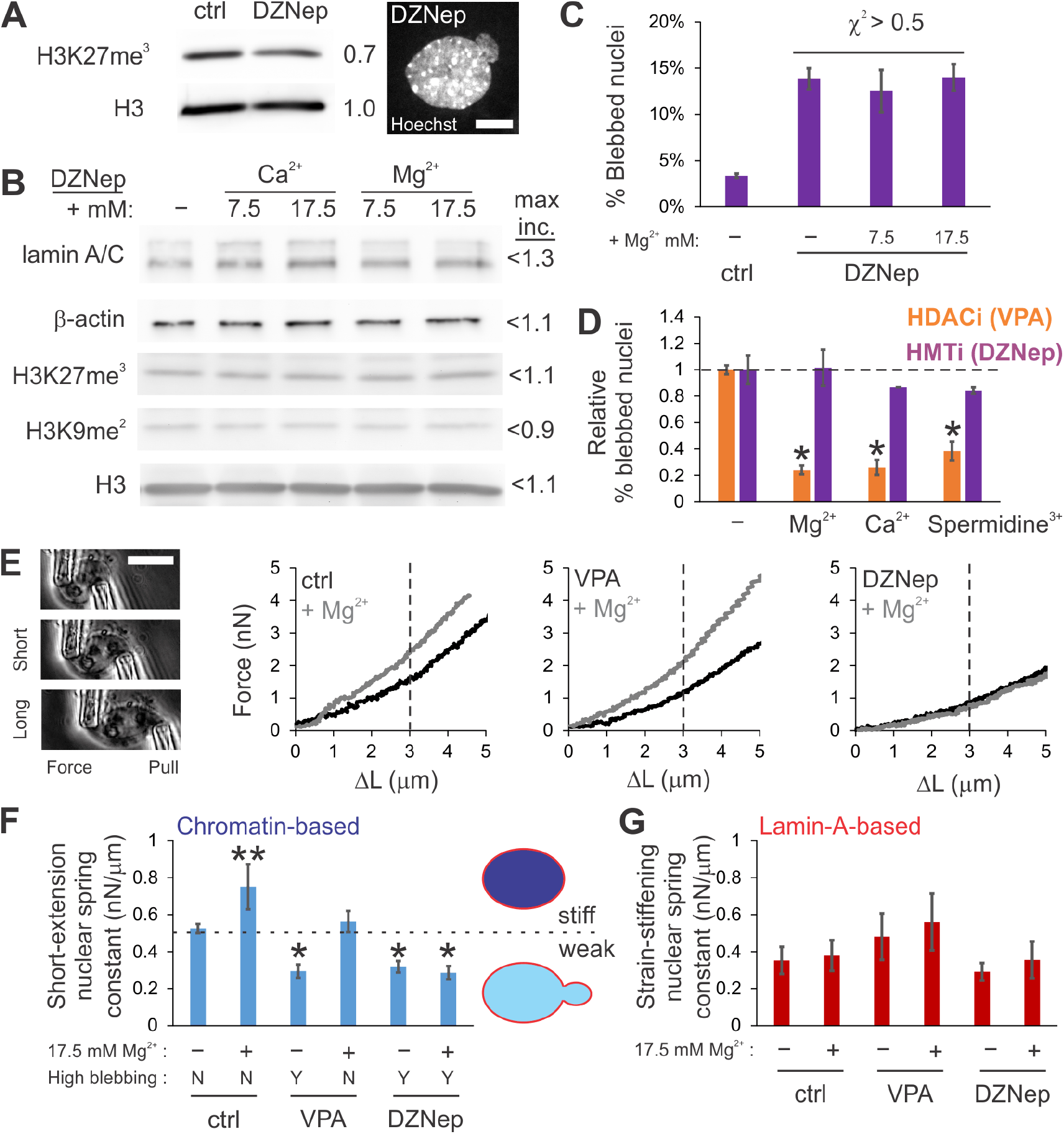
Increased extracellular divalent cations suppress nuclear blebbing via methyltransferase activity to increase heterochromatin and chromatin-based nuclear rigidity. (A) Western blots and an example of a blebbed nucleus after DZNep treatment. (B) Western blots of DZNep-treated MEF cells in standard or media supplemented with divalent cations for 24 hours. Maximum signal increase is listed. (C) Graph of percentage of DZNep-treated nuclei displaying a nuclear bleb and (D) relative nuclear blebbing for MEF VPA (orange) or MEF DZNep (purple) upon addition of extracellular cations (control scaled to 1). For all nuclear blebbing experiments, n = 3-5 experiments of 90-200 cells each, except for Ca^2+^ DZNep, which has n = 1 experiment. (E) Representative micromanipulation nuclear force measurement images and force vs. extension plots for control (ctrl), VPA, and DZNep pre-incubated cells in normal (black) or medium supplemented with 17.5 mM MgCl_2_ (gray). Vertical line separates mechanical regimes. (F,G) Graph of average nuclear spring constants measured by micromanipulation for (F) chromatin-based short extension (< 3 µm) and (G) lamin-A-based strain stiffening (n = 7-14, calculated as spring constant > 3 µm – spring constant < 3 µm). Bottom panel F denotes whether high nuclear blebbing is present as yes (Y) or no (N). Graphic depicts weaker short-extension nuclear spring constant due to decompacted chromatin that results in increased nuclear blebbing, while stronger nuclei do not display increased nuclear blebbing. Scale bar = 10 µm. Error bars represent standard error. Asterisks denote statistically significant differences and different numbers of asterisks are statistically significantly different from each other (p or χ^2^ < 0.05).

### Morphology rescue by heterochromatin formation occurs with an increase in the nuclear spring constant

Micromanipulation force measurements of isolated nuclei provide a way to measure the separate mechanical contributions of chromatin to short-extension stiffness and lamin A/C to strain stiffening at long extensions (Stephens et al., 2017b; Stephens et al., 2017a; Stephens et al., 2018). Nuclear spring constants were measured in MEF vimentin null (V-/-) cells for ease of nucleus isolation. MEF V-/-nuclei exhibit similar physical properties to wild type (Stephens et al., 2017a; Stephens et al., 2018), including nuclear bleb suppression via MgCl_2_ (Supplemental Figure 2G). Micromanipulation force measurements of isolated nuclei revealed that both VPA-and DZNep-treated nuclei, which display increased nuclear blebbing, were about 40% softer than control in the chromatin-dominated regime, but were similar to control in the lamin A/C-dominated regime (Figure 3E-G). This is consistent with previous reports that chromatin-based rigidity dictates nuclear morphology, independent of lamins (Stephens et al., 2018).

Because extracellular divalent cations increased heterochromatin levels, we predicted that cation treatment would increase chromatin-based nuclear spring constants. In all conditions, before performing force measurements, the medium was exchanged for fresh standard medium to avoid physicochemical chromatin condensation. Control MEF cells incubated for 24 hours in increased extracellular MgCl_2_ (17.5 mM) displayed a significant increase in the chromatin-based short-extension spring constant and no change in strain stiffening at long extensions (Figure 3E-G). Similar to control, nuclei from cells co-treated with VPA and additional extracellular MgCl_2_ exhibited a significant increase in the short-extension nuclear spring constant. The treatment rescued chromatin-based nuclear stiffness to a level comparable to control nuclei (Figure 3E and F) and decreased nuclear blebbing to near control levels (Figure 3D). Both lamin A/C mechanics and lamin A/C content were unchanged (Figure 3G; Figure 1D-F and 2D). This result supports our hypothesis that mechanosensitive ion channels can regulate histone modification state, which dictates chromatin-based nuclear rigidity and in turn, the nucleus’ ability to maintain normal shape.

We further tested this hypothesis by using DZNep as a downstream blockade to heterochromatin formation. Cells co-treated with DZNep and increased extracellular MgCl_2_ (17.5 mM) did not exhibit a rescued, stiff nuclear spring constant, but instead remained weak (gray line, Figure 3E DZNep). This demonstrates that the ability to increase the short-extension nuclear force response depends on heterochromatin formation via histone methyltransferase activity (Figure 3F, ctrl and VPA vs. DZNep). Consequently, the inability of DZNep-treated cells to form heterochromatin, even in response to extracellular ions, results in weaker nuclei that are susceptible to nuclear blebbing. This is likely due to heterochromatin’s mechanical contribution since global transcription inhibition does not disrupt divalent-cation-induced bleb suppression (Supplemental Figure 2H). Thus, in these experiments, mechanosensitive ion channels sensing extracellular ionic conditions cue histone methyltransferases to increase heterochromatin formation, which stiffens chromatin-based nuclear rigidity and maintains or rescues normal nuclear shape.

### Increased extracellular cation mechanotransduction rescues nuclear shape in model and patient cells of human diseases

We asked whether this mechanotransductive process can rescue nuclear morphology in diseased cells. A cellular model of a disease with abnormal nuclear morphology is ectopic expression of the mutant lamin A progerin, which is associated with the accelerated aging disease Hutchinson-Gilford progeria syndrome (HGPS) (Goldman et al., 2004; Butin-Israeli et al., 2012) and normal human aging (Rodriguez et al., 2009). As with many other known cases of abnormal nuclear morphology, GFP-progerin HeLa nuclei exhibited decreased heterochromatin levels, as measured by immunofluorescence and Western blots of H3K27me^3^ (Figure 4E, Supplemental Figure 3A, (Shumaker et al., 2006; McCord et al., 2013)). GFP-progerin-expressing HeLa cells do not display discrete nuclear blebs, but instead display abnormal overall nuclear shape, which can be quantified by a nuclear irregularity index (Figure 4, A and B, see Methods (Stephens et al., 2018)). Similar to observations in MEF and HT1080 cells, extracellular MgCl_2_ restored both nuclear elliptical shape and heterochromatin levels to near HeLa wild-type levels (Figure 4A-E, Supplemental Figure 3A). Rescue of nuclear shape was inhibited when HeLa GFP-progerin cells were co-treated with the mechanosensitive ion channel inhibitor GsMTx4 and extracellular MgCl_2_, confirming the dependence of rescue on mechanosensitive ion channels (Supplemental Figure 3B).

**Figure 4.**
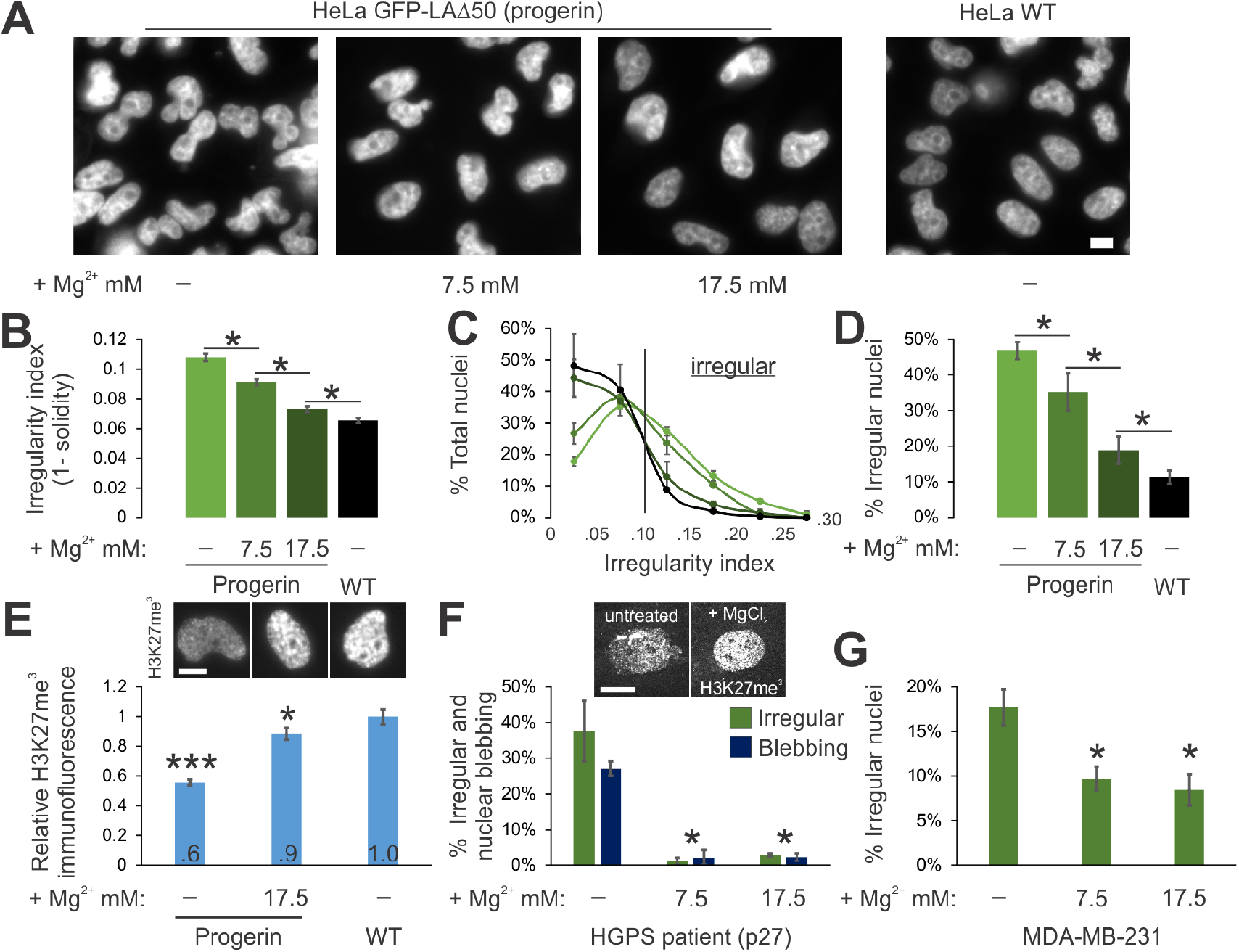
Increased extracellular divalent cations rescue nuclear shape in models and patient cells of human diseases. (A) Representative images of HeLa nuclei labeled with Hoechst. (B-D) Graphs of nuclear irregularity index showing (B) averages, (C) histograms, and (D) percentages of irregular nuclei (irregularity index > 0.10) for HeLa expressing GFP-progerin with various amounts of extracellular MgCl_2_ and wild type (WT) without additional MgCl_2_ (n = 2 experiments each with 160-281 cells). (E) Representative images and graph of relative immunofluorescence signal for facultative heterochromatin marker H3K27me^3^ for HeLa GFP-progerin in normal or MgCl_2_ supplemented media compared to HeLa WT (n > 100 cells). (F) Graph of percentages of nuclei that are irregularly shaped (green, irregularity index > 0.05) and display nuclear blebs (dark blue) for Hutchinson-Gilford progeria syndrome (HGPS) human fibroblast patient cells in normal or MgCl_2_ supplemented media (n > 30 cells, 2 experiments each). Representative images of facultative heterochromatin immunofluorescence are shown above the graph. (G) Graph of percentages of irregular nuclei (irregularity index > 0.10) in breast cancer model cell line MDA-MB-321 in normal or MgCl_2_ supplemented media (n = 4 experiments, each > 120 cells). Scale bar = 10 µm. Error bars represent standard error. Asterisks denote statistically significant differences (p or χ^2^ < 0.05), where different numbers of asterisks are also significantly different.

We next tested whether these observations hold for progeria (HGPS) patient cells and a breast cancer model cell line (MD-MBA-231). Progeria patient cells display abnormal nuclear shape and nuclear blebbing, but overall, patient cell nuclei are less irregular than HeLa cell nuclei. In progeria patient cells, increasing extracellular divalent cation concentration was sufficient to significantly decrease the percentage of irregular nuclei (here, given by index > 0.05) from 37% to 2% and nuclear blebbing from 29% to 2% (Figure 4F). Similarly, breast cancer model MD-MBA-321 cells treated with extracellular cations exhibited decreased mean nuclear irregularity index (from 0.07 without treatment to 0.06 with ions) and a decrease from 17% irregular nuclei (index > 0.10) without treatment to 8% irregular nuclei with additional cations (Figure 4G). Together, these data provide evidence that extracellular cues and chromatin contribute to nuclear organization and shape in a range of physiological conditions, including in a human aging syndrome and a model of cancer.

### Cells mechanotransduce extracellular signals to regulate chromatin state, nuclear rigidity, and nuclear morphology

We previously showed that changes to chromatin histone modification state modulate cell nuclear mechanics and shape (Stephens et al., 2017a; Banigan et al., 2017; Stephens et al., 2018). Here we have shown that such changes can be triggered by native cellular mechanotransduction via mechanosensitive ion channels. Alterations in the composition of the extracellular medium can induce heterochromatin formation by histone methyltransferases to modulate nuclear mechanics and morphology, without direct force application to cells. Our findings provide evidence that chromatin-based mechanics are tunable, physiologically relevant, and can be modulated via mechanotransduction across a range of cells. These effects are triggered by mechanosensitive ion channels embedded in the plasma membrane. Thus, direct stretching of cells (Le et al., 2016; Heo et al., 2016), cell-cell interactions, compression (Versaevel et al., 2012; Jain et al., 2013), ion composition (Figures 1 and 2), or charged proteins could all elicit this effect. The fact that our experimental perturbation involves only the extracellular environment highlights that this response is not dependent on distortion of the cell as a whole, but rather only on physiochemical signals received at and transduced through the cell membrane. Altogether, our results illuminate a general mechanism by which cells can mechanically adapt to different extracellular environments and maintain robust and normal nuclear morphology.

## Materials and Methods

### Cell growth

MEF, HT1080, and MDA-MB-231 cells were cultured in DMEM (Corning) complete with 1% Pen Strep (Fisher) and 10% Fetal Bovine Serum (FBS, HyClone) at 37°C and 5% CO_2_. HGPS patient cells were cultured in MEM complete medium with 1% Pen Strep and 15% FBS (Goldman et al., 2004). HeLa progerin (GFP-LAδ50) cells were grown in DMEM containing 1 mg/mL of G418 also at 37°C and 5% CO_2_. As outlined in (Taimen et al., 2009), progerin expression was induced via 2 *μ*g/mL deoxycycline treatment for 24 hours. Upon reaching confluency, cells were trypsinized, replated, and diluted into fresh media.

### Drug treatment

Cells were treated with histone deacetylase inhibitor valproic acid (VPA) at 2 mM for 16-24 hours to accumulate decompacted euchromatin, as previously reported in (Stephens et al., 2017a). To deplete compact heterochromatin, cells were treated with histone methyltransferase inhibitor (HMTi) 3-Deazaneplanocin-A (DZNep) at 0.5 μM for 16-24 hours (Miranda et al., 2009). To inhibit mechanosensitive ions channels, cells were treated with either GsMTx4 (2.5 μM) or GdCl_3_ (10 μM) for 16-24 hours.

### Immunofluorescence

Immunofluorescence experiments were conducted as described previously in Stephens et al. (Stephens et al., 2017a). Cells were seeded on cover glasses in six well plates and incubated at 37°C and 5% CO_2_. Once at 80-90% confluence, cells were fixed with 4% paraformaldehyde (Electron Microscopy Sciences) in PBS for 15 minutes at room temperature. Cells were then washed three times for 10 minutes each with PBS, permeabilized with 0.1% Triton X-100 (US Biological) in PBS for 15 minutes, and washed with 0.06% Tween 20 (US Biological) in PBS for 5 minutes followed by two more washes in PBS for 5 minutes each at room temperature. Cells were then blocked for one hour at room temperature using a blocking solution consisting of 10% goat serum (Sigma Aldrich Inc) in PBS. Primary antibodies were diluted in the blocking solution at the following concentrations: H3K27me^3^ 1:1,600 (C36B11, Cell Signaling), H3K9me^2-3^ 1:100 (6F12, Cell Signaling), lamin A/C 1:10,000 (Active Motif), and lamin B1 1:500 (ab16048, Abcam). After being incubated with the primary antibodies overnight at 4°C in the dark, cells were washed with PBS three times for 5 minutes each. Next, cells were incubated with anti-mouse or anti-rabbit Alexa 488 or 594 (Life Technologies, 2 mg/mL) fluorescent secondary antibodies diluted at 1:500 in blocking solution for one hour at room temperature in the dark. Cells were washed with 1 μg/mL Hoechst 33342 (Life Technologies) in PBS for 5 minutes and then washed three more times with PBS. Finally, cover slides were mounted onto microscope slides using ProLong Gold antifade reagent (Life Technologies) and allowed to dry overnight at room temperature.

### Western blots

Western blots were carried out as described previously (Stephens et al., 2017a; Stephens et al., 2018). Protein was extracted via whole cell lysates (Sigma) or histone extraction kit (Abcam). Protein was loaded and run in 4-12 % gradient SDS-polyacrylamide protein gels (LICOR) for an hour at 100 V. Gels were then transferred to nitrocellulose blotting membrane with 0.2 μm pores (GE Healthcare) via wet transfer for 2 hours at 100 V. The membrane was then washed three times in TBST for 5 minutes each before blocking in either 5% Non-fat Milk or BSA in TBST for an hour at room temperature. Primary antibody was diluted into 5% Milk or BSA, depending on company specifications, added to blotting membrane and allowed to shake and incubate overnight at 4°C. Primary antibodies were diluted in the blocking solution at the following concentrations: heterochromatin markers H3K27me^3^ 1:500 (Millipore) and H3K9me^2^ 1:500 (Millipore), euchromatin marker H3K9ac 1:2000 (Cell Signaling Technology), total histone control H3 1:2,000 (Cell Signaling Technology), lamin A/C 1:2,000 (Active Motif), and loading control β-actin 1:6,000 (Licor). The next day the membranes were washed four times with TBST before incubation in secondary antibodies conjugated to HRP (Millipore, 12-348 and 12-349) for 1 hour at room temperature. The membranes were again washed with TBST three times before chemiluminescence (PerkinElmer, Inc, NEL104001EA) and visualization via UVP imager. Quantification of Western blots was done in ImageJ.

### Imaging and analysis

Immunofluorescence images were acquired with an IX-70 Olympus wide field microscope using a 60X oil 1.4 NA Olympus objective with an Andor iXon3 EMCCD camera using Metamorph. Image stacks with a step size of 0.4 *μ*m were also acquired using a Yokogawa CSU-X1 spinning disk Leica confocal microscope with a 63X oil 1.4 NA objective and a Photometrics Evolve 512 Delta camera using Metamorph. Exposure times for DAPI, Rhodamine and FITC were between 50-600 ms. Images were saved with Metamorph and transferred to ImageJ for analysis. Nuclei were selected by ImageJ threshold detection in the brightest plane or drawn by hand around Hoechst fluorescence if nuclei were too close together. Background fluorescence was determined by quantifying a 30×30 pixels area with no cells. Average intensity values of the nuclei were acquired and the average background signal was subtracted using Excel. For comparisons between cell types or treatment conditions, relative intensities were reported as fold intensity relative to wild-type or untreated nuclei. Each nucleus was scored as blebbed if a protrusion larger than 1 µm in diameter was present, as detailed previously (Stephens et al., 2018). Solidity measurements were acquired in ImageJ following threshold detection. Statistical significance was determined for nuclear spring constants, immunofluorescence, and nuclear irregularity index measurements via the t-test. The chi-squared test was used to determine the statistical significance of changes in nuclear blebbing percentages.

### Live cell imaging

Cells were grown to the desired confluence in cell culture dishes containing glass coverslip bottoms (In Vitro Scientific). The dishes were treated with 1 μg/mL Hoechst 33342 (Life Technologies) for 10 minutes and then imaged on a wide field microscope as described above.

### Micromanipulation force measurement of an isolated nucleus

Micromanipulation force measurements were conducted as described previously in Stephens et al. (Stephens et al., 2017a). MEF vimentin null (V-/-) cells were used for their ease of nucleus isolation from a living cell and have similar nuclear force response to wild-type nuclei (Stephens et al., 2017a; Stephens et al., 2018). The nucleus was isolated by using small amounts of detergent (0.05% Triton X-100 in PBS) locally sprayed onto a living cell via a micropipette. This gentle lysis allows for a second micropipette to retrieve the nucleus from the cell via slight aspiration and non-specific adherence to the inside of the micropipette. Another micropipette was attached to the opposite end of the nucleus in a similar fashion. This “force” micropipette was pre-calibrated for its deflection spring constant, which is on the order of nN/µm. A custom computer program written in LabView was then run to move the “pull” micropipette and track the position of both the “pull” and “force” pipettes. The “pull” pipette was instructed to move 5 µm at 45 nm/sec. The program then tracked the distance between the pipettes to provide a measure of nucleus extension ∼3 µm. Tracking the distance that the “force” pipette moved/deflected multiplied by the pre-measured spring constant provides a calculation of force exerted. Calculations were done in Excel (Microsoft) to produce a force-extension plot from which the best-fit slope of the line provided a spring constant for the nucleus (nN/µm). Each nucleus was stretched 2-4 times to provide an accurate and reproducible measurement of the nuclear spring constant. Short-extension chromatin-based spring constants were measured from 0 – 3 µm and strain-stiffening lamin-A-based spring constants were measured as the difference between the short (0-3 µm) and long (3-6 µm) spring constants.

### ICP-MS analysis of cellular ion contents

MEF cells were grown to 50-60% confluency in 225-cm^2^ flasks, then changed into 25 mL of fresh growth medium only, growth medium containing 7.5 mM MgCl_2_, and growth medium containing 17.5 mM MgCl_2_, respectively. These cells were further incubated for 24 h at 37°C and 5% CO_2,_ and then harvested through trypsinization. Cell suspensions were centrifuged and aspirated to remove supernatant, and then 10 mL of fresh growth medium containing 100 µM Gd-DOTA (gadolinium-tetraazacyclododecane tetraacetic acid) was added. Cell mixtures were mixed for 5 minutes by gentle finger tapping and then centrifuged and aspirated to carefully remove supernatant. Cell pellet samples and drops of background 100 µM Gd-DOTA media were dried and then digested in 150 µL 67% nitric acid (Sigma, Trace grade). After digestion, 3000 µL MQ-H_2_O (the final nitric acid concentration will be 3% (wt)) was added to the samples. The 10-and 100-times diluted samples were further prepared by adding 3% nitric acid in MQ-H_2_O. All undiluted and diluted samples then ready for inductively coupled plasma mass spectrometry (ICP-MS) analysis. Thermo iCAP Q, Quantitative Bioelement Imaging Center of Northwestern University was used to determine total metal contents using one set of trace element calibration standards (200, 100, 40, 20, 10, 5, and 0.1 ppb). Sample element concentrations (ppb) were determined in Kinetic Energy Discrimination (KED) mode with 10% H_2_/He gas to minimize polyatomic interferences. ICP-MS data is analyzed via Microsoft Excel 2012. Extracellular element content was removed by using Gd-DOTA as tracer to subtract extracellular medium background since Gd-DOTA species remain outside of cells (Aime et al., 2002). Thus, cellular Mg, Ca and P element contents were calculated by subtracting the extracellular element content from the total (cellular + extracellular) element content. The molar ratios of cellular Mg and Ca, content relative to cellular phosphorus content (Mg/P and Ca/P) were used to normalize cellular metal content for different experiments because this normalization avoids inherent deviations from cell number and volume measurements for each experiment.

## Acknowledgements

We thank Yixian Zheng for providing us with MEF LB1-/-cells (Shimi et al., 2015). We thank Aykut Erbaş, Sumitabha Brahmachari, and Ronald Biggs for helpful discussions. We thank the Quantitative Bio-element Imaging Center (QBIC) at Northwestern University for ICP-MS analysis. A.D.S. is supported by Pathway to Independence Award NIHGMS K99GM123195 and wassupportedby NRSApostdoctoralfellowshipF32GM112422.A.D.S.,P.Z.L,E.J.B., and J.F.M. are supported by NSF Grants DMR-1206868 and MCB-1022117, and by NIH Grants GM105847, CA193419, and via subcontract to DK107980. S.A.A. and R.D.G. are supported by NIH GM106023, CA 193419 and Progeria Research Foundation PRF 2013-51. H.C. and T.V.O are supported by Physical Science – Oncology Center, NCI grant U54CA193419. L.M.A. and V.B. are supported by NIH grants R01CA200064, R01CA155284, NSF grant CBET-1240416, and Lungevity Foundation. This work was funded by the Chicago Biomedical Consortium with support from the Searle Funds at the Chicago Community Trust through a Postdoctoral Fellowship to A.D.S.

## Author contributions

A.D.S lead conceptualization, writing, and supervision of the paper while helping do data and analysis; P.Z.L lead data acquisition and analysis while helping with conceptualization and writing; V.K. and H.C. did data acquisition and analysis; L.M.A did conceptualization, V.B, T.O. S.A.A. and R.G helped with conceptualization and resources, E.J.B. lead conceptualization equally with A.D.S and helped with writing; J.F.M. did conceptualization, writing, supervision, and resources.

## Supplemental Figures

**Supplemental Figure 1.**
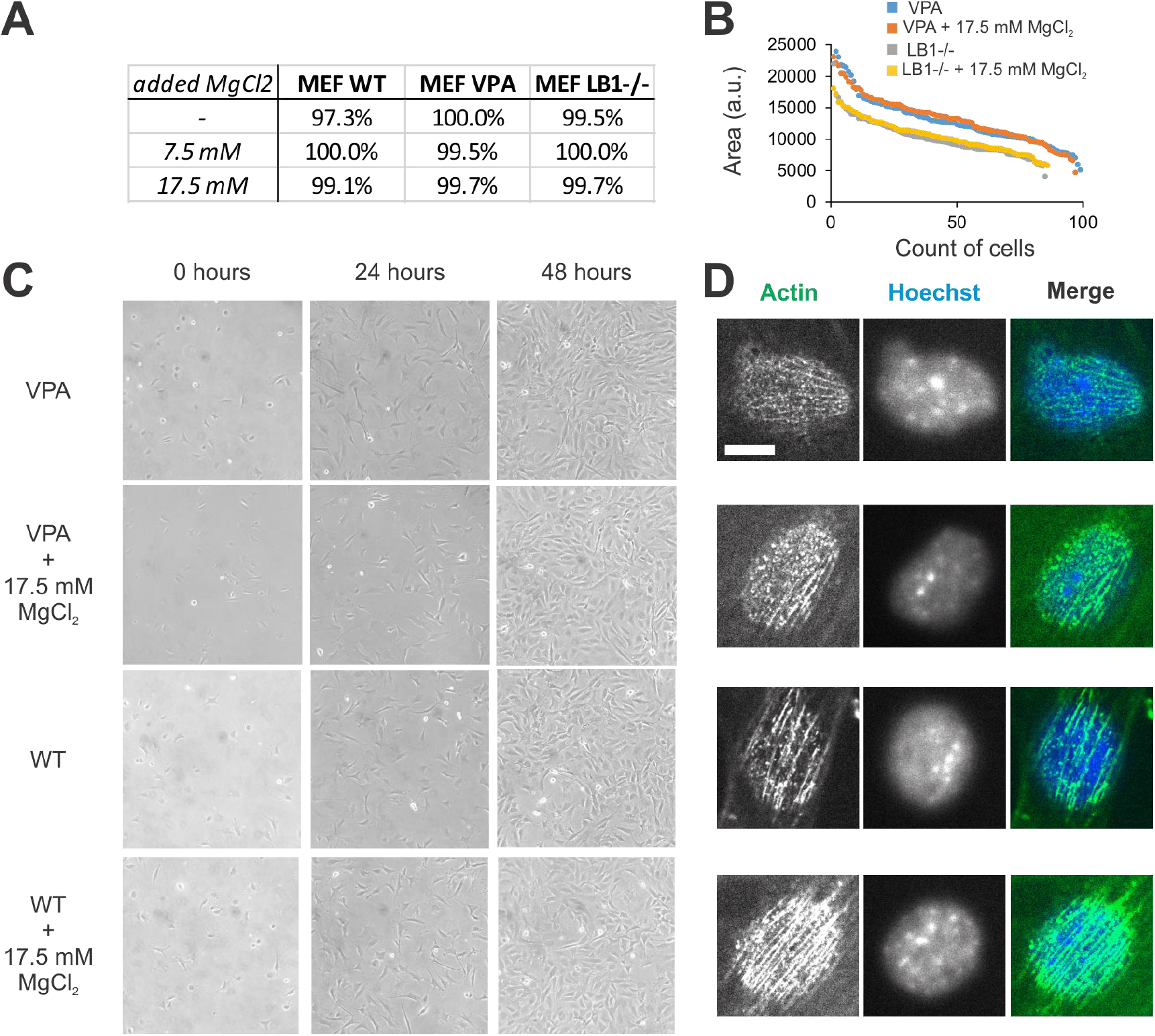
Increased extracellular divalent cations do not alter normal cell growth or characteristics. (A) Viability measured by the absence of propidium iodide staining, which marks dead cells (n = 200 – 330). MEF cells treated with VPA or null for lamin B1 (LB1-/-) in normal or MgCl2 supplemented media display similar (B-D) nuclear size, (C) cell confluency and growth, and (D) actin cables running over the nucleus. Scale bar = 10 µm.

**Supplemental Figure 2.**
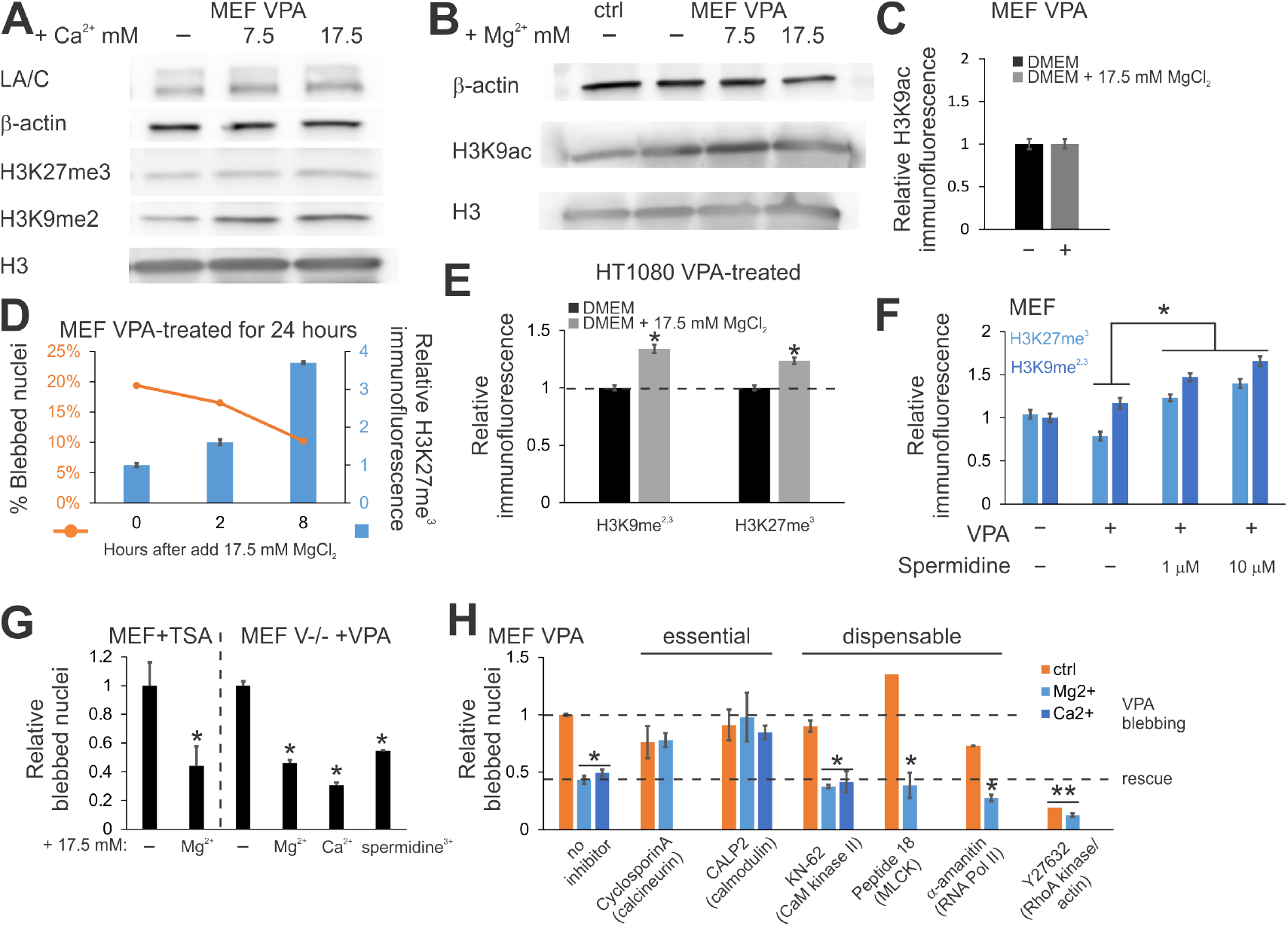
The extracellular amount of divalent or polyamine cations alters heterochromatin levels and nuclear blebbing over hours, but does not alter euchromatin levels. (A) Western blot of MEF VPA plus increased extracellular CaCl_2_. (B) Western blot and (C) IF of euchromatin marker H3K9ac in MEF VPA-treated cells with added MgCl_2_. (D) MEF nuclei treated with VPA for 24 hours, and then continued treatment in VPA with increased extracellular MgCl_2_ 17.5 mM for 0, 2, and 8 hours were measured for H3K27me^3^ immunofluorescence signal and nuclear blebbing percentage. Relative immunofluorescence signal of heterochromatin markers in (E) HT1080 cells treated with VPA and extracellular MgCl_2_ or (F) MEF cells treated with VPA and spermidine. (G) Relative nuclear blebbing in MEF TSA-treated or MEF V-/-VPA-treated without or with increased extracellular divalent cations (n= 2-4 experiments each > 100 cells). (H) Relative nuclear blebbing in MEF VPA-treated cells co-treated with inhibitors Cyclosporin A 5 µM, CALP2 20 µM, KN-62 8µM, Peptide 18 10 µM, α-amanitin 10 µM, and Y-27632 15 µM for 24 hours. Single asterisks in H denote decreased nuclear blebbing relative to control (ctrl, orange bar) and two asterisks denotes both control and divalent cation decreased relative to no inhibitor. Error bars represent standard error. Asterisks denote statistically significant differences (p < 0.01 or χ = 0.01), where different numbers of asterisks are also significantly different.

**Supplemental Figure 3.**
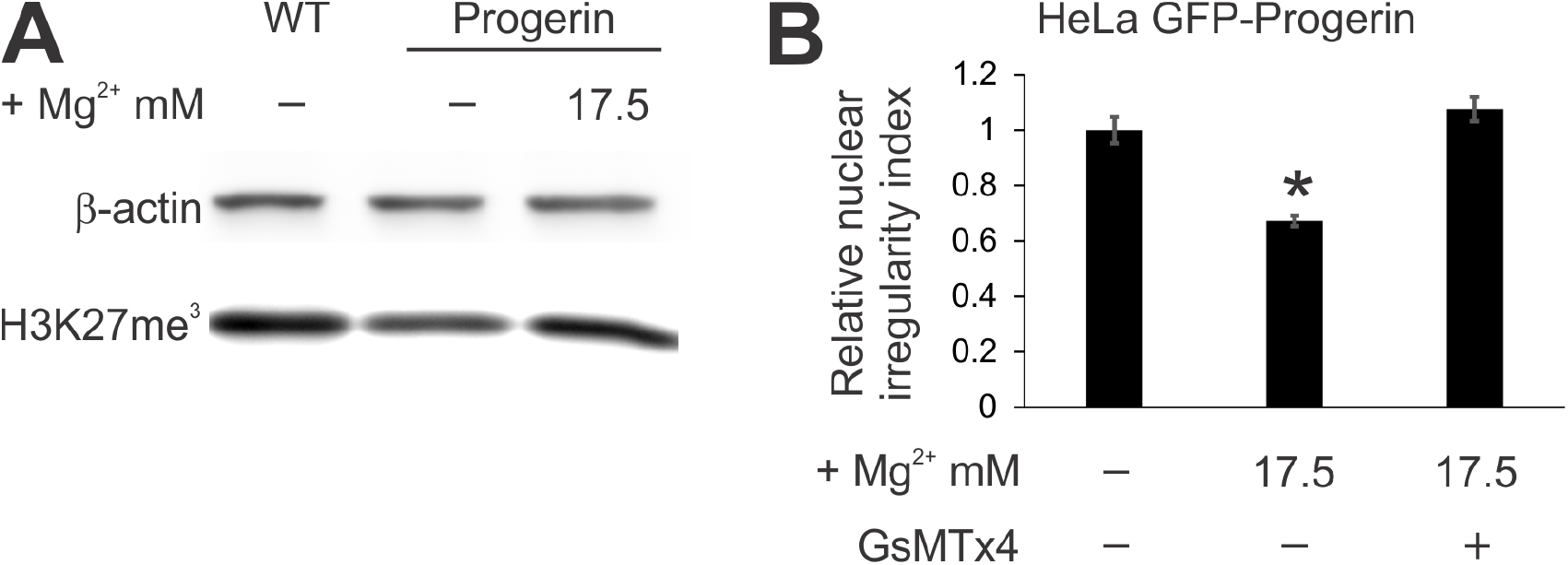
HeLa GFP-progerin expressing cells display extracellular MgCl_2_-induced heterochromatin increase and rescued nuclear morphology dependent on mechanosensitive ion channels. (A) Western blot of H3K27me^3^ in increased extracellular MgCl_2_. (B) Relative nuclear blebbing in HeLa GFP-progerin treated with just 17.5 mM extracellular MgCl_2_ and co-treated with GsMTx4 mechanosensitive channel inhibitor. Error bars represent standard error. Asterisks denote statistically significant differences (p < 0.01), where different numbers of asterisks are also significantly different.

